# Gamma-Protocadherins regulate filopodia self-recognition and dynamics to drive dendrite self-avoidance

**DOI:** 10.1101/2022.11.23.517768

**Authors:** Samantha Ing-Esteves, Julie L. Lefebvre

## Abstract

Neurons form cell type-specific morphologies that are shaped by molecular cues and their cellular events governing dendrite growth. One growth rule is distributing dendrites uniformly within a neuron’s territory by avoiding sibling or ‘self’ branches. In mammalian neurons, dendrite self-avoidance is regulated by the clustered Protocadherins (cPcdhs), a large family of recognition molecules. Genetic and molecular studies suggest that the cPcdhs mediate homophilic recognition and repulsion between self-dendrites but this model has not been tested through direct investigation of self-avoidance during development. Here we performed live imaging and 4D quantifications of dendrite morphogenesis to define the cPcdh-dependent mechanisms of self-avoidance. We focused on the mouse retinal starburst amacrine cell (SAC), which requires the gamma-Pcdhs (*Pcdhgs*) and self/non-self recognition to establish a stereotypic radial morphology while permitting dendritic interactions with neighboring SACs. Through morphogenesis, SACs extend a transient population of dynamic filopodia that fill the growing arbor and contact nearby self-dendrites. Compared to non-self-contacting filopodia, self-contacting events have longer lifetimes and a subset persists as filopodia bridges. In the absence of the *Pcdhgs*, non-self-contacting filopodia dynamics are unaffected but self-contact-induced retractions are significantly diminished. Filopodia bridges accumulate, leading to the bundling of dendritic processes and disruption to the arbor shape. By tracking dendrite self-avoidance in real-time, our findings demonstrate that the γ-Pcdhs selectively mediate contact-induced retractions upon filopodia self-recognition. Our results also illustrate how self-avoidance shapes the stochastic and space-filling behaviors of filopodia for robust dendritic pattern formation in mammalian neurons.

**HIGHLIGHTS:**

- Dendrite self-avoidance proceeds through interstitial filopodia and contact-induced retractions between sibling processes.
- Self-contacting filopodia exhibit longer lifetimes and a subset of contacts persist.
- Pcdhgs selectively regulate self-contact-induced retractions.
- Loss of *Pcdhgs* and filopodia self-avoidance disrupts dendritic arbor shape.

## INTRODUCTION

Morphological diversity of neurons is a critical feature of nervous system organization and function. Each neuron type forms dendritic and axonal arborizations with characteristic branch density and distribution (Lefebvre and Marocha 2020; Hoersting and Schmucker 2021). These wiring patterns develop from a protracted but highly dynamic period of neurite outgrowth, stabilization, and elimination. While numerous cell surface receptors involved in dendrite morphogenesis have been identified (Lefebvre 2021; Zang et al. 2021), we have a limited understanding of how the signals guide neurite dynamics to govern arbor patterning.

One growth rule for branch patterning is self-avoidance, where sibling branches from the same cell recognize and repel each other to prevent overlap (Grueber and Sagasti 2010; Kramer and Kuwada 1983; Lefebvre et al. 2015). Although self-avoidance is widely important for neuronal growth, the phenomenon is most apparent and studied in cells with stereotypic and two-dimensional arborizations, such as sensory neurons that innervate the epidermis in invertebrates (Gan and Macagno 1995; Grueber et al. 2003; Smith et al. 2010). In mammals, several neuron types form non-crossing arbors, including retinal and cerebellar Purkinje cells, and sensory neurons that project to the olfactory bulb and skin (Montague and Friedlander 1991; Lefebvre et al. 2012; Fujishima et al. 2012; Mountoufaris et al. 2017; Kuehn et al. 2019). A particularly striking example of self-avoidance is the radial arborization of the retinal starburst amacrine cell (SAC). SACs uniformly distribute their sibling or ‘self’ dendrites, but also synapse and fasciculate extensively with ‘non-self’ dendrites of ∼25-50 neighboring SACs, indicating a requirement for self/non-self recognition. The exquisite SAC morphology and dendritic plexus are needed to organize synaptic connectivity and generate direction selectivity in retina (Kostadinov and Sanes 2015; Morrie and Feller 2018), underscoring the importance of self-avoidance patterning for circuit function.

Self-avoidance requires the coupling of neurite self-recognition and repulsion. At the molecular level, dendrite self-avoidance can be accomplished by homophilic recognition molecules and ligand-receptor pairs that mediate repellent interactions (Lefebvre 2021). In *C. elegans*, branch spacing of the PVD sensory neuron is mediated by repellent Netrin/Unc-40/Unc-5 signaling between contacting self-dendrites (Smith et al. 2012; Sundararajan et al. 2019). Cell-autonomous signaling by secreted Wntless also influences PVD self-avoidance, as do Slit2/Robo2 interactions for Purkinje dendrite self-avoidance (Liao et al. 2018; Gibson et al. 2014). In *Drosophila* larvae, self-avoidance of dendritic arborization (*da*) neurons requires *Dscam1*, a gene that produces >18,000 isoforms with homophilic binding properties to mediate cell-specific homotypic interactions for self/non-self discrimination (Matthews et al. 2007; Hughes et al. 2007; Soba et al. 2007). At the cellular level, self-avoidance in *da* and PVD neurons converge on a mechanism of contact-induced retraction, as captured in live-imaging studies (Smith et al. 2010; Smith et al. 2012; Matthews et al. 2007). Recent 4D analyses of *da* arborization dynamics revealed a surprisingly stochastic dendritic growth that is shaped by *Dscam1* and contact-induced retractions (Palavalli et al. 2021; Shree et al. 2022). Observing self-avoidance in real-time is crucial for defining the mechanisms, but quantitative live imaging approaches have not been extended to mammalian neurite self-avoidance.

In the mammalian CNS, neurite self-avoidance is regulated by the clustered Protocadherins (cPcdhs). The cPcdhs comprise ∼60 cadherin-related molecules that are encoded by three linked gene clusters, *Pcdh-alpha, Pcdh-beta*, and *Pcdh-gamma* (Wu and Maniatis 1999). Deletion of *Pcdhg* causes loss of self-avoidance in SACs and Purkinje cells, resulting in dendrite crossings and altered arbor territories (Lefebvre et al. 2012). The cPcdhs also regulate self-avoidance of olfactory sensory axons, and other aspects of neurite patterning and survival in the CNS (Wang et al. 2002; Lefebvre et al. 2008; Prasad et al. 2008; Garrett et al. 2012; Suo et al. 2012; Hasegawa et al. 2016; Mountoufaris et al. 2017; Chen et al. 2017; Katori et al. 2017; Carriere et al. 2020; Mancia Leon et al. 2020). The cPcdhs generate extensive recognition specificity through combinatorial expression and homophilic binding of isoforms, providing a molecular basis for neurite self/non-self discrimination in mammals (Esumi et al. 2005; Kaneko et al. 2006; Schreiner and Weiner 2010; Rubinstein et al. 2015; Nicoludis et al. 2016; Goodman et al. 2016; Goodman et al. 2022; Thu et al. 2014; Brasch et al. 2019).

Elucidation of cPcdh functions in dendrite self-avoidance come largely from genetic dissection of *cPcdhs* in SACs. The *Pcdhgs* are required during development and act cell-autonomously in SACs (Lefebvre et al. 2012). SACs are unaffected in single *Pcdha* or *Pcdhb* cluster mutants, but dendritic crossings are severely exacerbated when diversity is further reduced in *Pcdha; Pcdhg* mutants indicating functional cooperation between cPcdhs (Ing-Esteves et al. 2018). Despite access to cPcdh diversity, SACs only require a single *Pcdhg* isoform for self-avoidance (Lefebvre et al. 2012). However, multiple *Pcdhg* isoforms are needed to allow dendritic overlap and synapse formation between neighboring SACs, consistent with self/non-self discrimination (Lefebvre et al. 2012; Kostadinov and Sanes 2015). The *Pcdhgs* are important for SAC circuit function by preventing the formation of autapses and promoting complete dendritic coverage critical for direction selectivity (Kostadinov and Sanes 2015). These studies led to a model in which the gamma-Pcdhs (γ-Pcdhs) mediate self-avoidance through contact-mediated repulsion of self-neurites (Lefebvre et al. 2012). However, this model was limited to inferences from *cPcdh* phenotypes in fixed cells and assumptions from invertebrates.

Here we performed time-lapse imaging of SAC morphogenesis to define the cellular dynamics and γ-Pcdhs-dependent events that drive dendrite self-avoidance. We imaged wild-type and conditional *Pcdhg* mutant SACs in retinal explants, and obtained high-resolution quantifications of dendritic structures and dynamics across development. In contrast to mature SACs that display a radial dendritic arbor with little branch variation (Poleg-Polsky et al. 2018), young SACs elaborate an exuberance of fine processes that sample the growing territory. This dynamic population of interstitial filopodia contact nearby ‘self’ dendrites and undergo contact-induced retraction that is dependent on the γ-Pcdhs. Our 4D quantifications reveal distinct self-contacting dynamics and γ-Pcdh-selective retractions that together provide a new understanding of how self-recognition self-organizes dendritic growth to establish a robust arborization pattern.

## RESULTS

### SAC morphogenesis is marked by exuberant dendritic filopodia growth

Mature SACs display a dendritic arbor with a stereotyped geometry and branching frequency, and a non-overlapping arrangement of sibling branches (Lefebvre et al. 2012; Poleg-Polsky et al. 2018). To determine how the SAC dendritic arbor pattern is established, we sought to quantitatively describe dendrite morphologies through development. As SACs are the only cholinergic cell type in the mouse retina, we used *ChAT-Cre* mice and Cre-dependent vectors (Ivanova et al. 2010; Rossi et al. 2011) to visualize single SAC arbor with membrane-targeted fluorescent proteins (**Figure 1A**). Postnatal day (P) 0 to P4 SACs were labeled by crossing *ChAT-Cre* to the *Rosa-mTmG* Cre reporter (Muzumdar et al. 2007), taking advantage of the initially sparse Cre recombination. SACs at P6 and older stages were visualized by injecting Cre-dependent adeno-associated virus (AAV) vectors encoding with YFP- or mClover-CAAX into retinas of P0-P1 *ChAT-Cre* animals (Ing-Esteves and Lefebvre 2021).

**Figure 1:**
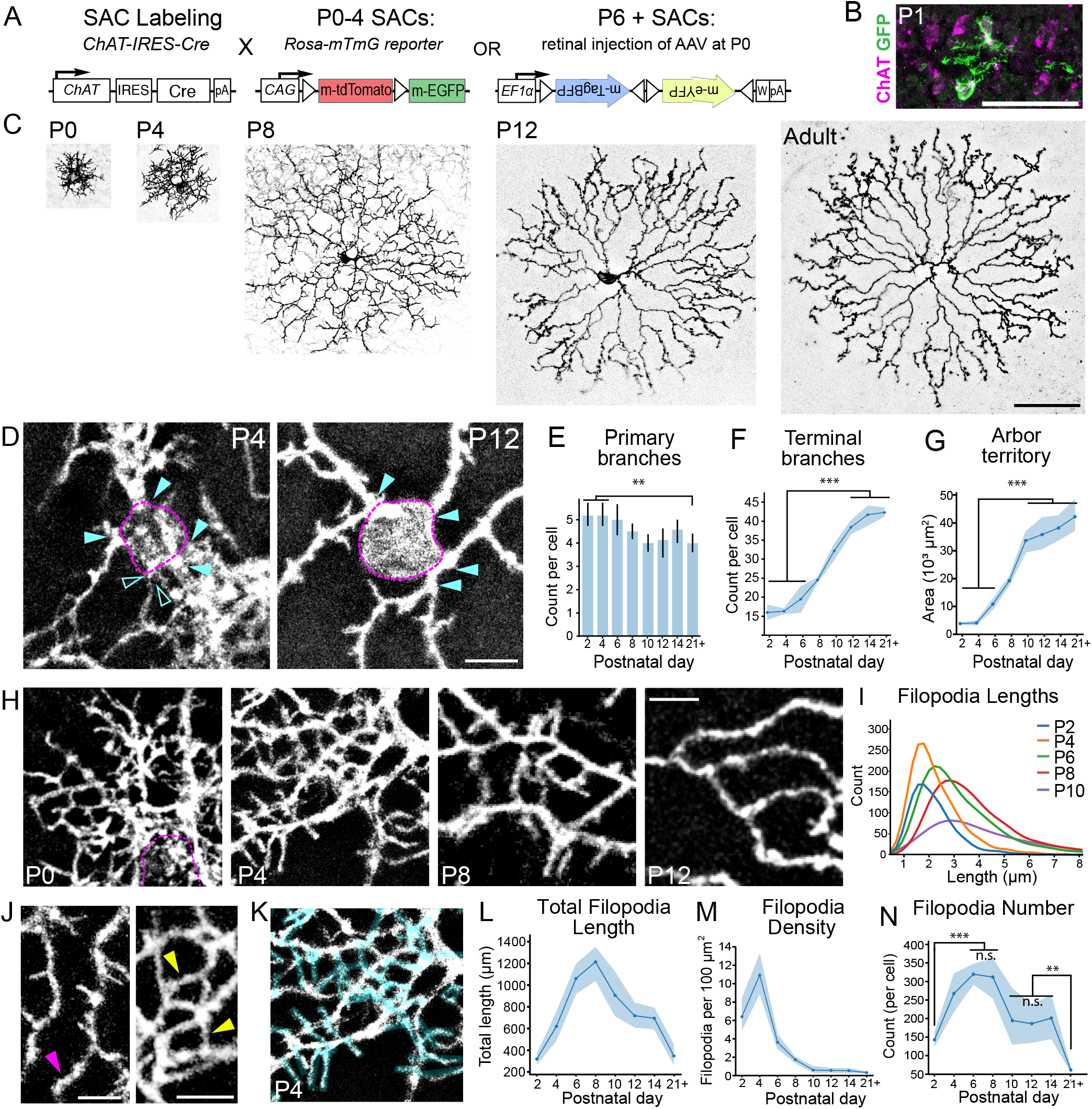
Developing Starburst Amacrine Cells (SACs) extend exuberant filopodia. **(A)** Schematic of SAC labeling strategy with membrane-targeted fluorophores in retina. *Chat-ires-Cre* mice were crossed to the *Rosa-mTmG* Cre-dependent reporter to visualize SACs at postnatal day (P) 0-4, or received intravitreal injection of Brainbow AAVs to visualize SACs at P6 and older. **(B)** Cross-section of *Chat-Cre; mTmG* retina at P1 shows two mGFP-labeled SACs (green) in the INL and GCL with dendrites ramifying within the IPL. SACs are immunolabeled for ChAT (magenta). **(C)** Single labeled SACs shown *en face* from P0-adult. Confocal images were captured in retinal flatmounts and shown as inverted maximum projections at the same scale. **(D)** P4 (left) and P12 SACs (right) showing the soma (magenta dotted line) and primary branches (cyan arrowheads). Unfilled cyan arrowheads depict a thin collateral that is typically not observed at maturity. **(E)** Number of primary branches per SAC through development. Bars show mean and 95% confidence intervals (CI). One-way ANOVA: F_7,59_ =3.78, p=0.0019, N= 6-11 cells from 3-8 animals per age. Post-hoc Tukey: ** p = 0.025; pairwise P8 to P21+, not significant. **(F)** Terminal branch points per SAC. Solid line shows mean and shaded lines are 95% CI. One-way ANOVA: F_7,45_=107.68, p<0.0001, N=6-9 cells from 3-7 animals per age. Post-hoc Tukey: ***p<0.0001. **(G)** Arborization area are quantified by convex hull area. Solid line shows mean and shaded lines are 95% confidence intervals (CI). One-way ANOVA: F_7,4_ =50.86, p<0.0001, N = 6-9 cells from 3-7 animals per age. Post-hoc Tukey: ***p<0.0001. **(H)** High magnification of SAC arbors depicting filopodia extension and refinement through development. **(I)** Filopodia lengths from P2 to P10, obtained from 6-9 cells from 3-7 animals per age. **(J)** Interstitial filopodia extend orthogonally from dendritic shafts (right, magenta arrowhead) and are distinctive from branch tip filopodia (left, yellow arrowhead). **(K)** Interstitial filopodia in P4 SAC from (H) are depicted with cyan overlay. **(L)** Total interstitial filopodia length per SAC from P2-maturity. **(M)** Number of interstitial filopodia per SAC from P2-maturity. One-way ANOVA: F_7,47_=12.15, p<0.0001. Post-hoc Tukey: ***p<0.002, **p<0.02. **(N)** Filopodia density. One-way ANOVA: F_7,47_=47.86, p<0.0001. Post-hoc Tukey: pairwise P8 to P21+ p=n.s.; ***p<0.001. Line plots in F, G, L-N are means with 95% CI, obtained from 6-9 cells from 3-7 animals per age. Scale bars are 50 μm in B, C; 5 μm in D, H, J.

We generated a developmental series of single-labeled SACs at two-day intervals spanning P0-P14 and at maturity, and quantified multiple features of dendritic arbor growth. Newborn SACs migrate to the ganglion cell layer or inner neuroblast layers during late embryonic stages, and extend dendrites within the nascent inner plexiform layer (Voinescu et al. 2009; Ray et al. 2018). By P1, SAC dendrites stratify and form the ON or OFF sublayers, which are visible in cross-sections (**Figure 1B**) (Ray et al. 2018; Stacy and Wong 2003). When viewed *on face* in retinal wholemount preparations, neonatal SACs display complex dendritic branching and short, filopodia protrusions (**Figure 1C**). Between P4 and P12, SAC arbors grow significantly in size and refine the filopodial processes. The characteristic self-avoidant pattern is apparent by P12. Postnatal SACs extend on average 5 primary dendritic branches, including a thin collateral branch emanating from the soma that is largely absent by P8-10 (**Figure 1D, E**). The number of terminal processes per SAC increases over postnatal development and peaks at P12-P14, similar to arbor area (**Figure 1F, G**).

One striking feature of developing SACs is the exuberance of short, filopodia-like processes within proximal and distal regions of the dendritic arbor (**Figure 1H**). In particular, a high density of interstitial processes project orthogonally from dendritic shafts, resembling interstitial filopodia described in mouse cortical dendritogenesis (Portera-Cailliau et al. 2003). We grouped these protrusions as ‘‘interstitial filopodia’ as they were similar in appearance and short, with mean lengths of 2.3 μm at P2-4 and 4.7 μm by P10, and similar in appearance (**Figure 1I**). Many filopodia appear to contact adjacent sibling or ‘self’ filopodia, forming looped structures and a webbed arbor pattern (**Figure 1H, 1J, 1K**). We quantified the interstitial filopodia of complete SAC arbors through the postnatal stages. Interestingly, the total number of filopodia per SAC peaks between P4-P8 (319 ± 33 filopodia per cell at P6) and then declines, despite the increasing territories of the cells (**Figure 1L, M)**. As such, filopodia density peaks at P4 and then declines sharply after P8 (**Figure 1N)**. Together, this analysis reveals that SAC arbor growth includes an exuberant elaboration and then elimination of filopodia during the first two postnatal weeks.

### Developing SAC dendrites form self-contacting filopodia bridges

Many interstitial filopodia appear to contact ‘self’ by forming bridges to nearby sibling filopodia or dendritic branches (**Figure 1H, J**, and **Figure 2A**). Self-contacts are rarely observed between distal tips of dendritic branches. Rather, they are predominantly located between filopodia tips and dendrite shafts within proximal regions of the arbor (**Figure 2A, B**). We confirmed the bridge structures within single confocal z planes (**Figure 2B,C**).

**Figure 2:**
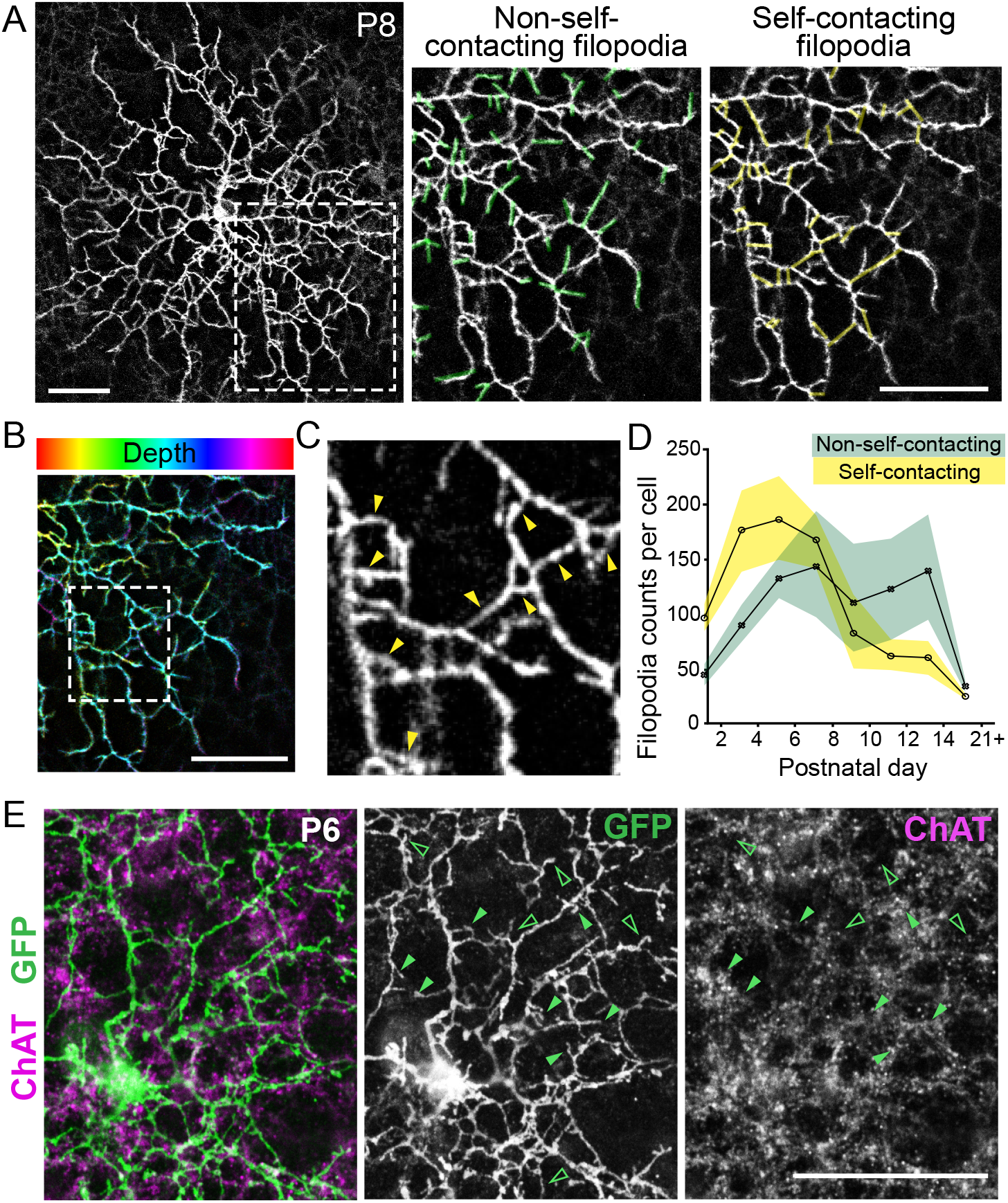
SAC dendrites form self-contacting filopodia bridges. **(A)** MembraneYFP-labeled SAC at P8. In enlarged views, yellow overlay denotes filopodia that contact isoneuronal or ‘self’ processes (right) and green overlays denote filopodia that do not self-contact (middle). **(B)** Maximum z-projection of total SAC volume with spectrum code depicting positions in z depth. **(C)** Single optical section showing boxed region from (B) confirms the positioning of self-contacts (yellow arrows). **(D)** Quantifications of non-self- (green) and self-contacting (yellow) filopodia through development. Line plots show mean with shaded regions as 95% CI. N = 6-9 cells from 3-7 animals per age. Two-way ANOVA show interaction with age and filopodia type, F_7,1_ =6.37, p<0.0001. Post-hoc Tukey test for P4, p = 0.052. **(E)** Colocalization of SAC dendritic branches and filopodia (green) with SAC dendritic plexus labeled with anti-ChAT (magenta). Many non-contacting filopodia overlap with ChAT+ SAC plexus (green filled arrowhead) but some do not (unfilled green arrows). Scale bars are 25 μm.

To compare the prevalence of the self-contacting to non-self-contacting filopodia, we divided the interstitial filopodia within SAC 3D image volumes using the following criteria: 1) ‘non-self-contacting’ filopodia have terminal endings (**Figure 2A**, center); 2) ‘self-contacting’ filopodia are bridge-like protrusions that form a continuous fluorescent signal with an adjacent sibling branch or filopodia (**Figure 2A**, right). The number of non-self-contacting filopodia increases until P6 and subsequently plateau. Self-contacting filopodia are most abundant from P4 to P8 (one-way ANOVA *p* < 0.0001; P2 vs P4, *p* = 0.002; P8 vs P10, *p* = 0.0014, post-hoc Tukey), and are more abundant than non-contacting filopodia at earlier stages (**Figure 2D**).

Dendrites of neighboring SACs fasciculate and form dendro-dendritic synapses with each other, forming a dense dendritic plexus that can be visualized by ChAT immunolabeling (Lefebvre et al. 2012; Kostadinov and Sanes 2015). We observed that many but not all self-contacting and non-self-contacting filopodia overlay with the ChAT plexus at P6 and P8 (**Figure 2E**), indicating that both populations are largely, but not entirely, in contact with processes of nearby SACs. Together, the quantifications of static SACs reveal a prevalence of dendritic filopodia that contact ‘self’ and form loops during morphogenesis.

### Developmental elaboration of dendritic filopodia is stochastic

We also found that developing SACs exhibit considerable variation in the number of filopodia extensions, with coefficients of variation ranging from 24% to 51% from P4-P8 (%CV = σ/mean*100; Table S1). The developmental variability in filopodia number contrasts with the relatively consistent number of dendritic branching and terminal segments reported for adult SACs (Poleg-Polsky et al. 2018; CV = 7% and 5%, respectively).The filopodia variability from SAC to SAC suggests that stochastic filopodia growth is a feature of SAC morphogenesis.

### Time-lapse imaging of developing SAC arbors reveals self-contacting filopodia dynamics as central structures in self-avoidance

We wondered whether the self-contacting filopodia are the substrates for Pcdh-gamma (Pcdh-γ)-dependent dendrite self-avoidance. To determine the role of self-contacting filopodia during dendrite morphogenesis, we investigated filopodia behaviors using a live confocal imaging pipeline we developed to capture SACs residing in the ganglion cell layer in retina flat mounts (Ing-Esteves and Lefebvre 2021). We visualized fluorescently-labeled SAC processes at P2, P4, P6, and P8. To limit the potential effects of phototoxicity on dendrite dynamics, images were acquired every 2 minutes over a 30-65 minute duration **(Supplemental Videos S1-5)**. The time-lapsed videos revealed highly dynamic and exploratory filopodia behaviors, including protrusions that iteratively extend and retract, and some of which contact self and others that do not. A smaller proportion of filopodia persisted through the end of the imaging period (**Figure 3A**). For each video, we quantified four populations of filopodia behaviors within selected ROIs: self-contacting filopodia that (i) retract or (ii) persist; and non-self-contacting filopodia that (iii) retract, or (iv) persist (**Figure 3B, Video S5)**. Filopodia extension events were highest at P2 and P4 (**Figure 3C, top**), consistent with the peak in filopodia density observed in complete SAC arbors (**Figure 1N**). The majority of filopodia retract, illustrating the high degree of filopodia motility (**Figure 3A, 3C)**.

**Figure 3:**
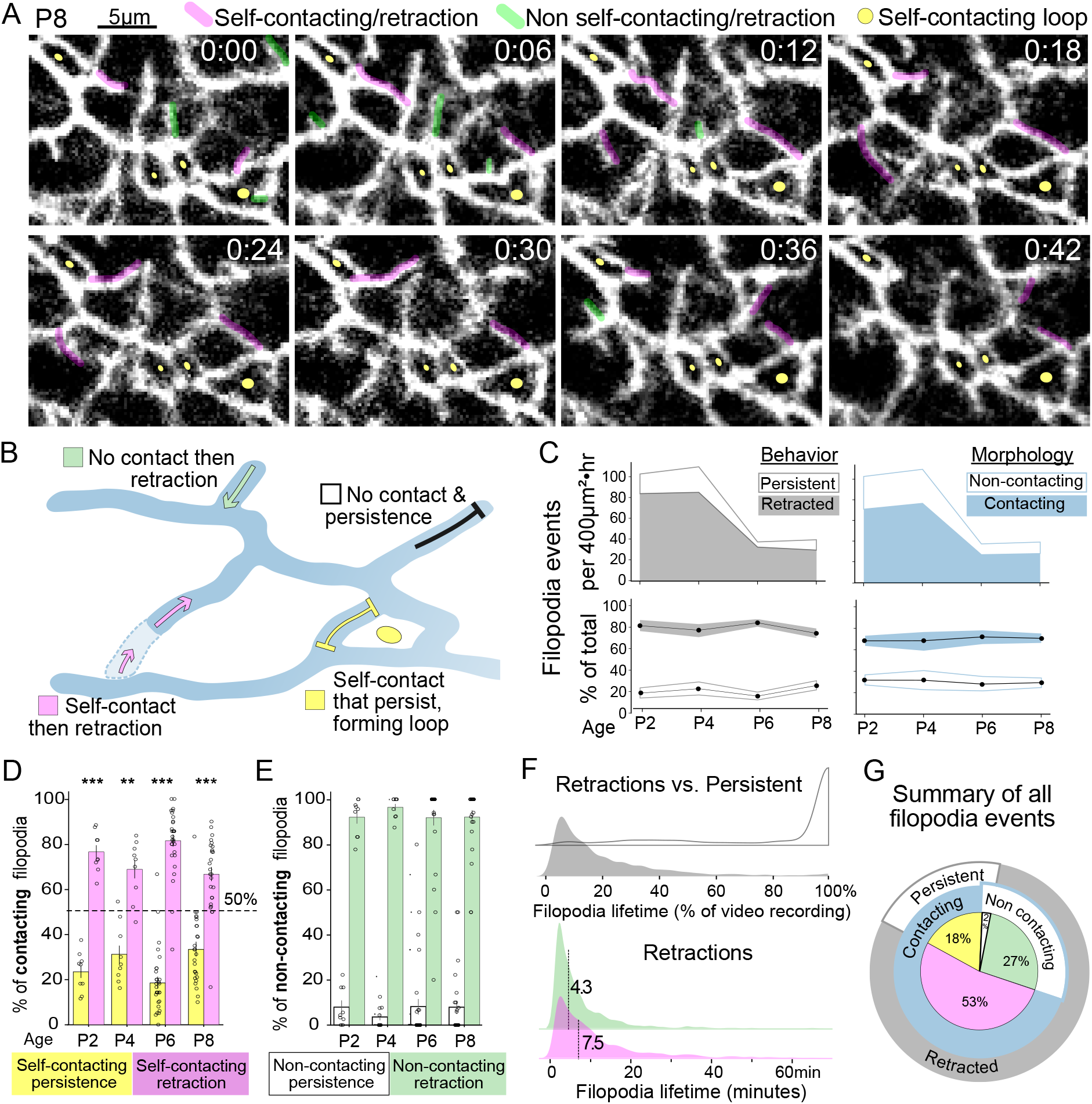
Time-lapse analysis of SAC dendrite morphogenesis reveals self-contact-induced retractions. **(A)** Still frames from Video S5 of P8 SAC. Frames were acquired 2 minutes but 6 minute intervals are shown here. Colored overlays denote filopodia behaviors. **(B)** Schematic of filopodia behaviors: Self-contacting filopodia that retract (magenta) or persist through the end of the video (yellow), and non-contacting filopodia that retract (green) or persist (white). **(C)** Quantifications of persistent and retracted filopodia events (left), and self-contacting and non self-contacting filopodia (right), reported as events per 400μm²•hr (top) and as percent of total obser-vations (bottom). Data are N= 9-28 ROIs from 3-7 cells, and 3-5 animals per age. **(D)** Quantifications of self-contacting filopodia that retract (pink) or stabilize (yellow). 1-sample t-tests were performed to compare to a 50% retraction rate. ***, p <0.0001 for all ages, except P4 **, p = 0.003. **(E)** The majority of non-self-contacting filopodia retract (green) rather than persist (white). Distribution bar plots in D, E, show mean +/- SEM from 9-28 ROIs from 3-7 cells, and 3-5 animals per age. **(F)** Distribution of filopodia lifetimes, which is the interval between filopodia extension and retraction. *Top*, Lifetimes of retracting (grey) versus persistent filopodia populations (white), plotted as a percentage of total video duration. *Bottom*, Self-contacting filopodia population (pink) has a longer median duration (7.5 minutes, N=1462 events) than non contacting filopodia (4.3 minutes, N=734 events; 2-sample Kolmogorov-Smirnov test: KS-statistic=0.72, p-value<0.0001). Data are pooled from 72 ROIs from P2-P8. **(G)** Pie chart summarizing all filopodia events observed in P2, P4, P6, P8 video datasets.

To determine if self-contacting filopodia dynamics differ from the non-self-contacting population, we compared their prevalence and motility. Across P2 to P8, most filopodia extensions result in a self-contact (**Figure 3C**). While the majority of self-contacting filopodia retract, a notable fraction is immotile and persists as loops through the imaging period (**Figure 3D**). By contrast, non-contacting filopodia rarely persist (**Figure 3E**). Finally, we compared their filopodial lifetimes, which are on the order of minutes. Interestingly, the self-contacting retraction population has a significantly longer median duration compared to the non-contacting median durations of (7.5 minutes vs 4.3 minutes, P2-P8; *p* < 0.0001, K-S test; **Figure 3F**). Of the persistent self-contacting filopodia, most are present for the entire duration of the video (**Figure 3F**). We could not quantify the eventual retractions of the longer lasting structures beyond the imaging durations.

Through live observation and quantifications of dendritic structures, we establish that SAC dendrite-self avoidance comprises the following cellular events: (1) SACs extend a transient population of exploratory, interstitial filopodia that fill the arbor and contact sibling or ‘self’ processes; (2) the majority of filopodia extensions result in self-contacts and subsequent retractions; (3) self-contacting filopodia exhibit longer filopodia lifetimes, suggesting a self-recognition event; and (4) a subset of filopodia self-contacts persist for long durations, a behavior seldom observed among non-self-contacting filopodia (**Figure 3G**). As longitudinal imaging of SACs over days was not possible, we could not track how filopodia dynamics influence the stabilization of dendritic branches, which are increasingly apparent by P8. However, from these short-term dynamic descriptions, we propose that dendrite self-avoidance is an iterative process of stochastic filopodia growth and contact-induced retraction that is dependent on self-recognition.

### SACs lacking the *Pcdhgs* accumulate filopodia self-contacts

Having established that dendrites in wild-type SACs self-avoid through filopodia extensions and retractions, we next sought to identify the specific functions of the gamma-Pcdhs (*Pcdhgs*). Visualization of GFP-tagged endogenous γ-Pcdhs in cultured SACs isolated from *Pcdhg*^*fcon3*^ mice confirmed the distribution of γ-Pcdhs along SAC dendritic shafts and protrusions, including at sites of self-contacts (**Figure 4A**). We next examined the effects of *Pcdhg* deletion on SAC morphogenesis by crossing *ChAT-IRES-Cre* to *Pcdhg*^*fcon3*^ mice (*Pcdhg*^*fcon3/fcon3*^; *ChAT-IRES-Cre*, herein referred to as *Pcdhg*^*SACko*^ for SAC-specific knockout, **Figure 4B)** (Lefebvre et al. 2008; Prasad et al. 2008). *Pcdhg*^*SACko*^ SACs have profoundly altered arbors, with dendritic bundles and gaps forming within the territory as early as P4 (**Figure 4C**), similar to pan-retinal *Pcdhg* mutant SACs (Lefebvre et al. 2012). *Pcdhg*^*SACko*^ SACs also extend an exuberance of interstitial filopodia but peak in total number later than wild-type SAC (355 ±110 filopodia at P8 compared to 319±33 at P6 in control SACs) (**Figure 4D, 4E)**. By P14, the filopodia are eliminated in *Pcdhg*^*SACko*^ SACs to a similar extent as in controls (**Figure 4D, 4E**). Therefore, postnatal *Pcdhg*^*SACko*^ SACs follow a program of dendritic growth and elimination similar to wild-type SACs.

**Figure 4:**
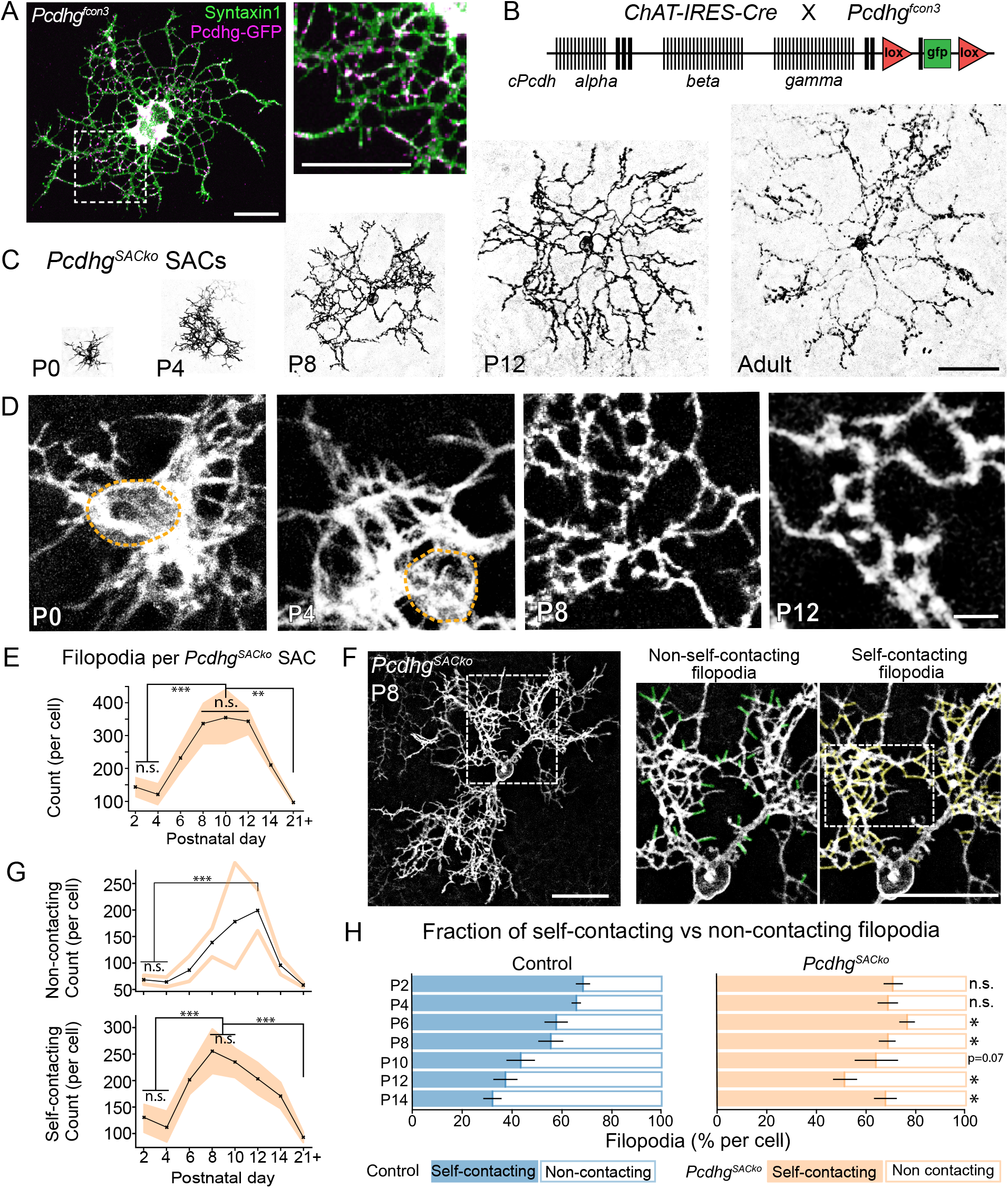
Self-contacting filopodia accumulate in *Pcdhg* mutant SACs. **(A)** SAC dissociated from *Pcdhg*^*fcon3*^ retinas and cultured *in vitro* displays Pcdhg-GFP proteins (anti-GFP, magenta) at dendritic tips, bridges and shafts (anti-syntaxin, green). **(B)** Schematic of the *cPcdh* locus with loxP-targeted *Pcdhg*^*fcon3*^. In the absence of Cre, the *Pcdhg*^*fcon3*^ allele produces Pcdhg-GFP fusion proteins due to Gfp fused to the C-terminus of constant exon 3. **(C)** Inverted fluorescent images of *Pcdhg*^*SACko*^ SACs from P0-adult, shown at the same scale. Arbor territory gaps are visible by P4. Accumulation of filopodia self-contacts is apparent at P8 but eliminated by maturity, while dendrite crossings are apparent by P12. **(D)** High magnification of *Pcdhg*^*SACko*^ SAC arbors depicting filopodia self-contacts. **(E)** Number of interstitial filopodia per *Pcdhg*^*SACko*^ SAC from P2-maturity. Solid line shows mean with 95% CI. One-way ANOVA F_7,45_= 11.022, p<0.0001, N = 6-8 cells, 3-6 animals per age. **p<0.003, post-hoc Tukey test for pairwise comparisons. **(F)** *Pcdhg*^*SACko*^ SAC at P8 extends non-self-contacting (green overlay, middle panel) and an excess of self-contacting filopodia bridges (yellow, right panel). **(G)** Quantifications of non-self-contacting (top, outlined) and self-contacting filopodia (bottom, orange) per SAC arbor. Line plots show mean with shaded regions as 95% CI. N = 6-9 cells from 3-7 animals per age. **(H)** Proportion of self-contacting (filled) versus non-contacting (unfilled) filopodia, shown as a percentage of total interstitial filopodia counts per cell for control (blue) and *Pcdhg*^*SACko*^ (orange). Error bars represent standard error of the mean. Self-contacting events for control and *Pcdhg*^*SACko*^ for each age were compared by two-sample t-test. *p<0.05, **p<0.007, ***p<0.0001. Scale bars are 25 μm in A, F; 50 μm in C, 5 μm in D.

*Pcdhg*^*SACko*^ SACs extend both self-contacting and non-self-contacting filopodia but display a striking accumulation of bridges in areas with higher branch density (**Figure 4F)**. *Pcdhg*^*SACko*^ SACs initially extend similar numbers of self-contacting filopodia as in controls but then accumulate significantly more self-contacts by P6 (**Figure 4G; 4H**, *p* < 0.05). While control SACs extend a higher prevalence of non-contacting filopodia by P12 (one sample t-test, *p* < 0.002), *Pcdhg*^*SACko*^ SACs maintain a higher proportion of self-contacting filopodia throughout development (**Figure 4H;** one sample t-test, *p* < 0.01 for all stages except P10, *p* = 0.17 ). We also noted that self-contacting filopodia appear to fasciculate, which is resolved by P12 (**Figure 4C)**. Therefore in the absence of the *Pcdhgs*, SACs extend a similar growth of dendritic filopodia but most maintain self-contacts. By maturity, *Pcdhg* SACs eliminate the filopodial protrusions, similar to wild-type SACs, but dendritic branch crossings are apparent (**Figure 4C, E, G**).

### The γ-Pcdhs selectively regulate self-contact-induced retractions

To determine the role of γ-Pcdhs in filopodia behaviors and self-contacting events, we analyzed time-lapse videos of SACs in *Pcdhg*^*SACko*^ mutant retinas (**Figure 5A, B, Supplemental Video S6-10**). We observed that non-self-contacting filopodia along *Pcdhg*^*SACko*^ dendrites emerge and retract similarly to control SACs but that self-contacting filopodia have visibly reduced motility (**Figure 5A**). Similar to wild-type SACs, filopodia events in *Pcdhg*^*SACko*^ SACs were categorized as one of four behaviors: (i) self-contacting filopodia that retract; (ii) or persist; and non-self-contacting filopodia that (iii) retract, or (iv) persist (**Figure 5B)**. There were no significant differences in the number of filopodia extensions nor in the number of self-contacting and non-self-contacting events sampled in *Pcdhg*^*SACko*^ ROIs compared to control ROIs (**Figure 5C-E**). The non-self-contacting filopodia behaviors are similar in *Pcdhg*^*SACko*^ and control SACs, with the exception of a small decrease in the rate of non-self-contacting retractions at P8 (14.8% in *Pcdhg*^*SACko*^ SACs vs 27.0% in WT SAC, two-way ANOVA, F(1,3)= 0.92, *p* =0.43; Games-Howell post-hoc pairwise comparison, *p =* 0.038) (**Figure 5F**). Moreover, there is no change in the occurrence of non-self-contacting filopodia that persist, indicating that *Pcdhg* loss does not enhance the stabilization of these filopodia (**Figure 5F)**. These results show that the general mechanisms for filopodia extensions and retractions are not disrupted in *Pcdhg* mutant SACs. We conclude that the *Pcdhgs* are dispensable for non-self-contacting filopodia dynamics.

**Figure 5:**
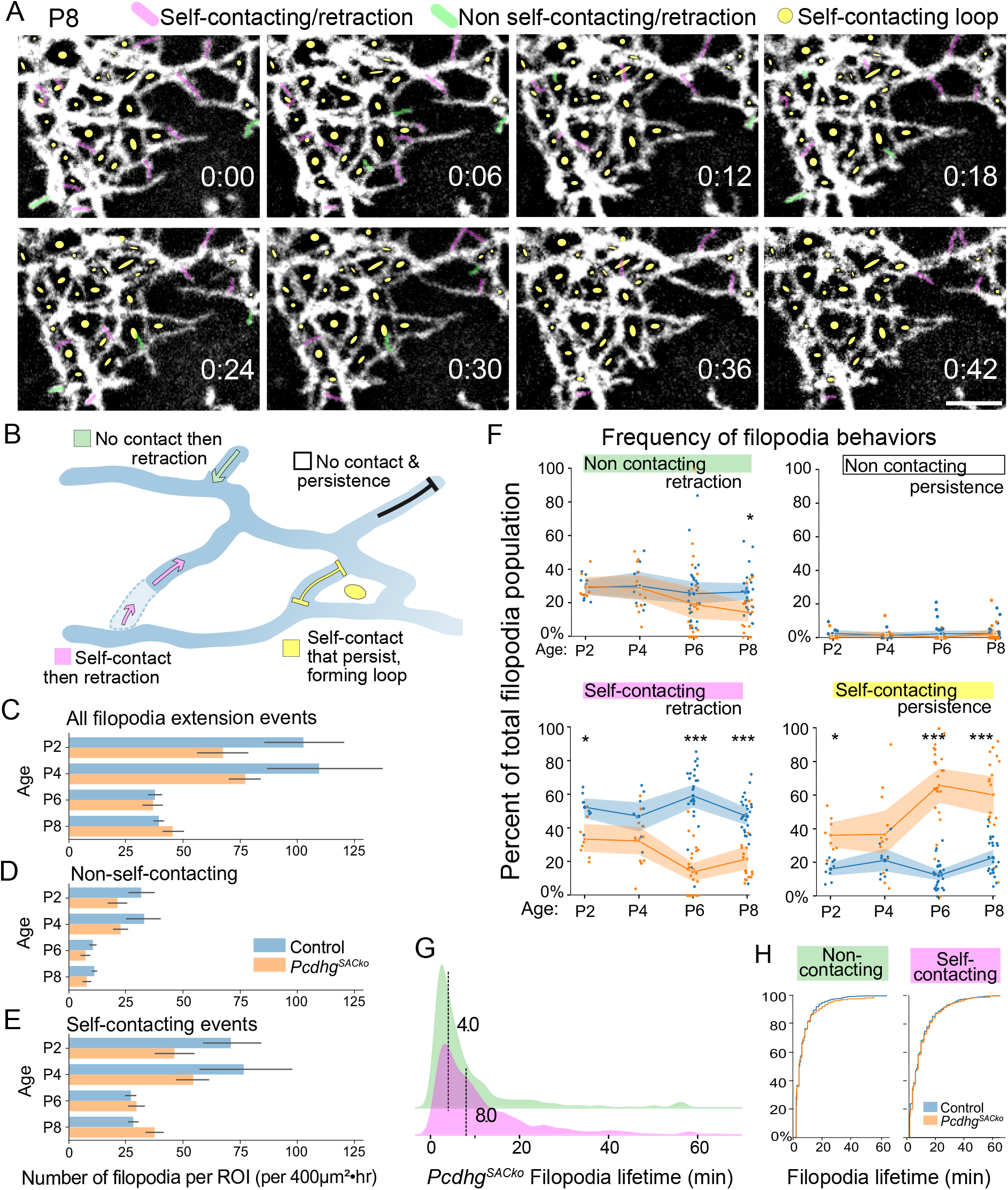
Dynamic measurements of *Pcdhg* SAC filopodia reveal that non-contacting events are unaffected but self-contacting retractions are reduced. **(A)** Still frames show 6 minute intervals of P8 *Pcdhg*^*SACko*^ SAC from Video S10, which was imaged every two minutes. Filopodia behaviors include self-contact-induced retractions (magenta); non-self-contacting and retraction (green); and self-contacting filopodia that are immotile to the end of the video (yellow dots).Scale bar, 5 μm. **(B)** Schematic of filopodia behaviors: Self-contacting filopodia that retract (magenta) or persist (yellow), and non-contacting filopodia that retract (green) or persist (white). **(C)** Quantifications of total filopodia extensions per ROI (400^2^μm•hour), imaged in control (blue) and *Pcdhg*^*SACko*^ SACs (orange) from P2 to P8. Bars show mean with SEM. Two-way ANOVA; Genotype [F_1,131_ =7.7, p = 0.006]; Genotype*Age [F_1,3_ =3.6, p = 0.02]. Games-Howell post-hoc pairwise comparisons for control versus *Pcdhg*^*SACko*^ mutants for each age group are not significant. N= 8-26 ROIS, 3-7 cells, 3-6 animals per age per genotype. **(D-E)** Data from C are subdivided into non-self-contacting and self-contacting filopodia events. Two-way ANOVA non-self-contacting filopodia events (D): Genotype [F_1,131_ =11.4, p = 0.001]; Genotype*Age [F_1,3_ =1.0, p = 0.4]. Two-way ANOVA for self-contacting filopodia events (E). Genotype [F_1,131_ =3.7, p = 0.05]; Genotype*Age [F_1,3_ =3.6, p = 0.02]. Games-Howell post-hoc pairwise comparisons are not significant. **(F)** Filopodia motility over development in control (blue) and *Pcdhg*^*SACko*^ SACs (orange). Each filopodia behavior is represented as a percentage of the total filopodia events. Line plots show means with shaded regions as 95% CI, from N= 8-26 ROIs, 3-7 cells, 3-6 animals per age per genotype. Two-way ANOVAs were performed for each group, testing for interactions with genotype or age. Non-self-contacting retractions in control versus *Pcdhg*^*SACko*^ SACs; F =2.4, p =0.12. Games-Howell post-hoc pairwise comparison, P8: p = 0.038. Self-contacting retractions, control versus *Pcdhg*^*SACko*^: F_1,131_=86.7, p < 0.0001. Post-hoc comparisons, P2: p = 0.030; P6, P8, <0.0001. Self-contacting persistence in control versus *Pcdhg*^*SACko*^: F =87.4, p < 0.0001. Post-hoc comparisons, P2: p = 0.012; P6, P8, <0.0001. * p <0.05, *** p <0.0001. **(G)** Filopodia lifetimes in *Pcdhg*^*SACko*^ SACs show longer median duration for self-contacting/ retraction population (pink) (dashed lined: 8.0 minutes, N=675 events) compared to non-contacting/retractions (dashed lined: 4.0 minutes, N=568 events). Events are pooled from 63 ROIs from P2-P8, and represent. **(H)** Cumulative distribution plots comparing self-contacting (left) and non-self-contacting lifetimes (right) for control (blue, data from Fig. 3F) and *Pcdhg*^*SACko*^ SACs (orange, data from Fig. 5G). Kolmogorov-Smirnov test: non-self-contacting: KS-statistic = 0.098, p = 0.0042. Self-contacting: KS-statistic = 0.098, p =0.43.

The most striking effect of *Pcdhg* deletion is on self-contacting filopodia dynamics. Filopodia self-contacts that retract are significantly reduced in *Pcdhg*^*SACko*^ compared to control SACs (**Figure 5F**). There is a sharp increase in the proportion of self-contacting filopodia that persist in *Pcdhg*^*SACko*^ SACs, with a 30% increase from P4 to P6 (**Figure 5F;** 2-way ANOVA F_1,3_= 8.04, *p* < 0.0001; P4 to P6, *p* = 0.03, Games-Howell post-hoc pairwise comparison). The rise in persistent self-contacts is significantly higher in *Pcdhg*^*SACko*^ than in control, which maintains a low rate through postnatal development. Since similar numbers of self-contacting filopodia events were sampled within wild-type and *Pcdhg*^*SACko*^ SAC ROIs (**Figure 5E)**, these quantifications indicate that *Pcdhg* deletion specifically diminished the rate of self-contact-induced retractions. *Pcdhg* loss does not slow filopodia retractions, as the population of self-contacting *Pcdhg*^*SACko*^ filopodia that retracts has similar dynamics as those in wild-type SACs (**Figure 5G, H;** *Pcdhg*^*SACko*^ median lifetime of 10.5 minutes, N = 675 self-contacting/retracting filopodia events; versus wild-type median lifetime of 10.9 minutes, N = 1462 events; 2-sample Kolmogorov-Smirnov test: KS-statistic = 0.04, p-value = 0.43). Thus, the γ-Pcdhs do not regulate filopodia retraction dynamics *per se*, but rather the choice to retract or persist.

From the quantifications, we conclude that the γ-Pcdhs mediate dendrite self-avoidance through contact-induced retractions between sibling processes. The γ-Pcdhs are dispensable for non-self-contacting filopodia dynamics, despite significant overlap and potential interactions with neighboring SAC dendrites (**Figure 2E**; Ray et al. 2018). The selectivity for self-contacts further supports the model that the γ-Pcdhs induce retractions upon self-recognition from non-self.

### *Pcdhg*-dependent dendrite self-avoidance establishes the radial arbor morphology

Finally, we asked when SACs achieve their characteristic radial shape as a result of Pcdhg-dependent self-avoidance. We quantified the morphologies of wild-type and *Pcdhg*^*SACko*^ SACs through development using two measures to describe radial shape: circularity and polar distribution of branches (Ing-Esteves et al. 2018). Arbor circularity is the ratio between convex hull area and perimeter, where a perfect circle has a circularity of 1. Wild-type SAC arbors at P2 are noticeably irregular in shape but increase in circularity by P10 (**Figure 6A, B**). Developing *Pcdhg*^*SACko*^ SAC arbors also increase in circularity but are significantly lower than wild-type at P14 (**Figure 6A,C)**.

**Figure 6.**
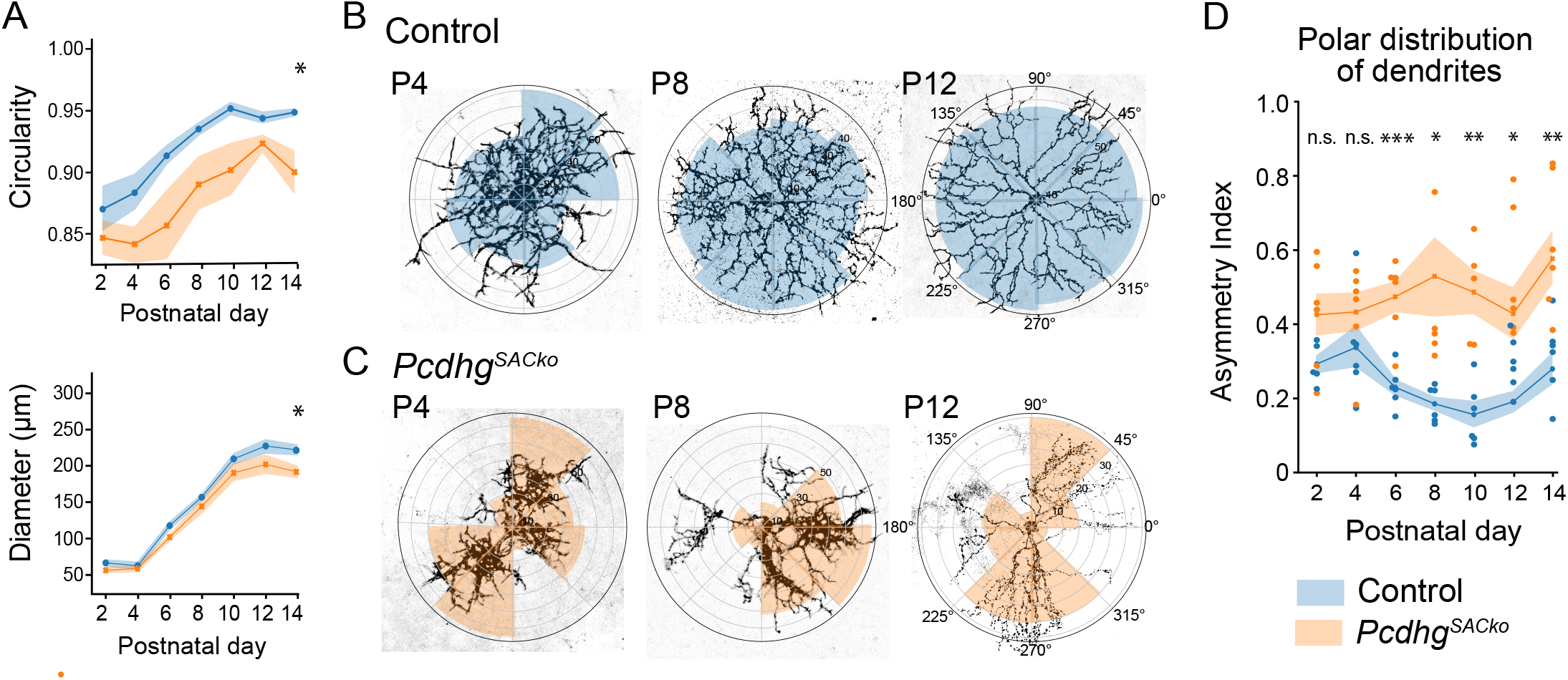
Postnatal *Pcdhg*^*SACko*^ SACs fail to develop symmetric dendritic arborizations. **(A)** Circularity (top) and diameters (bottom) of SAC arbors in control (blue) and *Pcdhg*^*SACko*^ (orange) retina through postnatal development. Circularity is the ratio between convex hull area and perimeter, where a circularity of 1 denotes a perfect circle. Line line plots show the mean with SEM, from N=6-9 cells, N=3-7 animals per age. P14: 2-sample t-test, *p<0.05. **(B)** Representative images of control SACs overlaid on their associated polar plots (blue) from which the polar distribution (or asymmetry) index is calculated. Shaded sectors depict the number of dendrite pixels from binarized images distributed within that arbor slice. The normalized standard deviation between these slices is the index value. **(C)***Pcdhg*^*SACko*^ SAC images and their polar plots (orange). **(D)** Polar distribution index of SAC dendritic distributions in control (blue) and *Pcdhg*^*SACko*^ (orange) retinas. Zero is a perfect radial value. Line line plots show the mean with SEM. Control and *Pcdhg*^*SACko*^ SACs compared by 2-sample t-tests: *p<0.05, **p<0.01, ***p<0.001.

To measure the distribution of branches, we calculated an asymmetry index that reports on the dendritic content divided across eight sectors of the SAC arbor (Ing-Esteves et al. 2018). As SAC morphogenesis proceeds, control cells establish a radial branch distribution (**Figure 6B, D)**. By contrast, *Pcdhg*^*SACko*^ SACs fasciculate filopodia and longer dendritic processes, and a significantly higher asymmetric index by P6 compared to control SACs (**Figure 6C, D)**. By P12, *Pcdhg*^*SACko*^ SACs eliminate the excess filopodia and self-contacts, as in wild-type, but then display dendrite crossings and bundling. Thus, the self-avoidance phenotypes in *Pcdhg* SACs do not simply result from branches growing over one another. Rather, the perturbed branch distribution and final arbor shape arise from the inability to retract self-contacts and the increased self-fasciculation during morphogenesis.

In conclusion, our analyses of dendrite dynamics on the scale of minutes show that self-avoidance is mediated by transient, interstitial processes that sample the available territory through self-contact and γ-Pcdh-dependent retractions (**Figure 7**). Together with analyses of arbor morphogenesis over days, this study also illustrates the iterative, self-organizing features of dendrite self-avoidance that generate the characteristic, non-overlapping SAC dendrite morphology.

**Figure 7:**
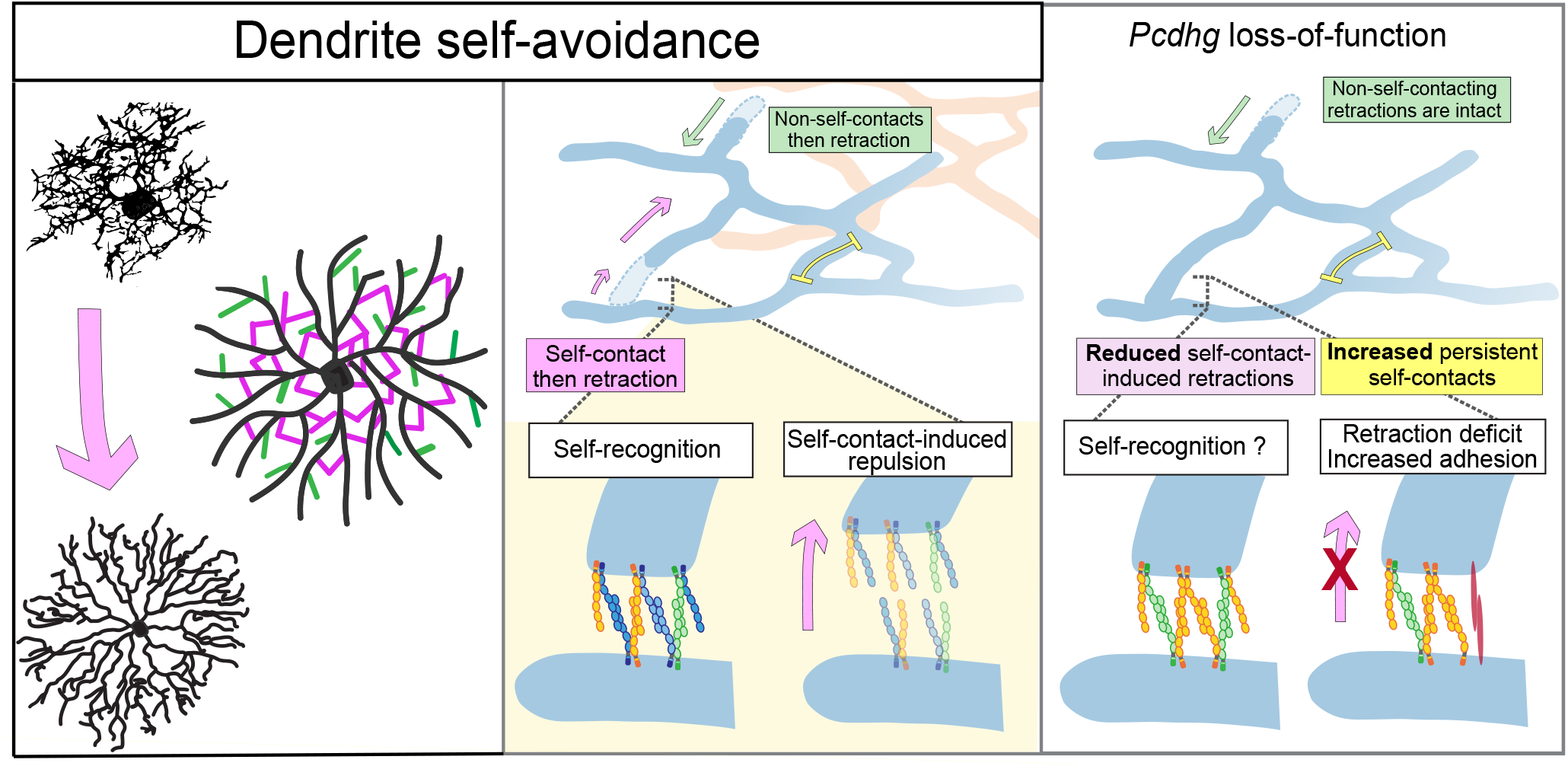
Model for Pcdhg-dependent dendrite self-avoidance and self/non-self descrimination. **Left**, Dendrite self-avoidance shapes the characteristic Starburst amacrine cell radial arborization (top, postnatal SAC; bottom mature SAC). Self-avoidance is driven by transient filopodia that iteratively self-sample (magenta) and retract through the γ-Pcdhs to generate a non-overlapping dendrite distribution. **Center top**, Dendrite self-avoidance is marked by self-contact-induced retractions between sibling filopodia processes (magenta arrow). Compared to non-contacting filopodia (green, which might overlap with neighboring SACs, orange), self-contacting filopodia exhibit longer lifetimes and a subset persist, forming bridges (yellow). **Center bottom**, the γ-Pcdhs mediate self-avoidance through a proposed model of homophilic binding of cPcdh isoforms to signal self-recognition and γ-Pcdh-dependent retraction (blue isoforms). **Right**, loss of *Pcdhg* disrupts self-contact-induced retractions. The accumulation of bridges suggest that isoneuronal interactions are preserved but self-recognition driven retractions are diminished.

## DISCUSSION

We previously identified the cPcdhs as molecular regulators of dendrite self-avoidance and required for non-overlapping dendritic coverage (Lefebvre et al. 2012; Ing-Esteves et al. 2018). In this study, we defined the cellular dynamics and *Pcdhg*-dependent events of dendrite self-avoidance through live observation and quantifications of dendritic growth. Dendrite self-avoidance couples space-filling filopodia growth with γ-Pcdh-dependent interactions between sibling processes to limit overlap. Remarkably, self-contacting (isoneuronal) filopodia interactions have longer lifetimes and their retraction rates are selectively diminished in the absence of γ-Pcdhs. By contrast, non-self-contacting events are unchanged in *Pcdhg* SACs, despite heteroneuronal dendritic interactions between neighboring SACs (Ray et al. 2018). Our results establish that the γ-Pcdhs regulate self-contact-induced retractions, and provide further evidence for self versus non-self discrimination (**Figure 7**). Morever, the self-contacts that persist for longer durations suggest a local decision to retract or persist, and that their dendritic loops further shape the self-avoidant pattern. We propose that self-avoidance is a negative feedback for stochastic dendritic growth through γ-Pcdh-driven self-recognition and retraction (**Figure 7**). The iterative self-sampling across the radial plane constrains branch growth along non-overlapping sectors. In this way, the self-organizing process driven by γ-Pcdh-mediated self/non-self discrimination and retraction produces a stereotyped, non-overlapping radial morphology.

### Contact-induced retraction is a conserved feature of mammalian dendrite self-avoidance

Our previous understanding of dendrite self-avoidance was limited to inferences of *cPcdh* phenotypes in fixed cells and from live imaging of PVD and *da* sensory neurons (Ing-Esteves et al. 2018; Lefebvre et al. 2012; Grueber and Sagasti 2010). The parallels between *Drosophila Dscam1* and mouse *Pcdhg* phenotypes, stochastic expression and binding properties also shaped the model that cPcdhs execute self-avoidance through homophilic repellent interactions between neurites arising from a single neuron (Zipursky and Sanes 2010; Lefebvre et al. 2012). Our analyses of SAC dynamics substantiates the cPcdh model and reveals important details of mammalian dendrite self-avoidance, including shared and distinct features from invertebrates. Similar to live observations of *C. elegans* PVD and *Drosophila da* dendrites (Matthews et al. 2007; Smith et al. 2010; Palavalli et al. 2021; Shree et al. 2022), Self-contacting filopodia have exploratory dynamics with extensions-retractions on the order of minutes, similar to the lifetimes of *Drosophila da* dendrites (Palavalli et al. 2021; Shree et al. 2022). Importantly, the core mechanism of contact-induced self-avoidance is conserved for mammalian neurons.

A limitation of this study is that the characterizations of filopodia contacts are limited to light microscopy. Given the diffraction limits, we cannot confirm that the opposed ‘contacting’ processes are in direct physical contact. We called self-contact events based on continuous fluorescent intensity, a criteria used in other recent studies (Palavalli et al. 2021; Shree et al. 2022; Han et al. 2012). We also limited calls to processes that abut and form ‘loops’ but do not pass over each other, as branch overlaps are typical of neurites in separate z-planes (Han et al. 2012; Kim et al. 2012). Self-contacts have been validated by electrophysiology in adult *Pcdhg* mutant SACs, as they form autapses because of diminished self-avoidance (Kostadinov and Sanes 2015). Whether postnatal self-contacts are functionally detectable has not been determined. Further experimentation using electron microscopy of genetically-labeled neurons, or super-resolution imaging of live cells could further resolve isoneuronal contacts in developing SACs.

### SAC dendrites self-avoid through interstitial filopodia

A notable feature of SAC dendrite self-avoidance are the interstitial filopodia, which are transient, space-filling structures and the main sites of self-contact and retraction. While these protrusions might simply be shorter dendrites or branchlets (Stürner et al. 2019), they resemble interstitial filopodia found along dendritic shafts within developing rodent pyramidal arbors and *Xenopus* tectal arbors (Portera-Cailliau et al. 2003; Dailey and Smith 1996; Hossain et al. 2012). In these cells, interstitial filopodia are precursors for spines or persist to become branches. Whether a subset of SAC interstitial filopodia stabilizes to form main dendritic branch is unknown and will require longer-term imaging and tools to track dendrite branch stabilization.

Our quantitative analyses indicate that the majority of interstitial filopodia form self-contacts. They are observed in proximal and in distal regions of the arbor, presumably in areas with a high density of sibling processes that increase the probability of short-range contacts. Thus, a compelling idea is that SACs extend space-filling filopodia to maximally sample the locations of sibling neurites and enhance the negative feedback needed for robust pattern formation. *Drosophila da* dendritic arborizations, which bear stereotyped morphologies, also exhibit a surprising degree of stochastic growth and elimination of processes during morphogenesis (Ferreira Castro et al. 2020; Palavalli et al. 2021; Shree et al. 2022). In Class 1 *da* arbors, the extension and *Dscam1*-mediated self-avoidance of transient processes contribute to branch stabilization and pattern formation. Space-filling dendritic extension is also observed in growing Class IV arbors, the most highly branched of the *da* neurons (Shree et al. 2022). Our work together with recent studies in invertebrates illustrate the importance of stochastic and space-filling interstitial growth in dendrite pattern formation.

Filopodia bridges and self-avoidance behaviors have been described in Purkinje cells (Sdrulla and Linden 2006). As in SACs, filopodia bridges are more apparent at younger stages and disappear as dendrites stabilize in a non-overlapping arrangement. Moreover, Purkinje dynamics captured *in vitro* exhibit motile filopodia and dendritic tips that display contact-dependent retractions from sibling and heteroneuronal dendrites (Fujishima et al. 2012). The Purkinje cells require both *Pcdhg* and *Pcdha* for dendrite self-avoidance (Ing-Esteves et al. 2018; Lefebvre et al. 2012) but the cPcdh-dependent dynamics remain to be investigated. Slit2/Robo2 and Sema6A/Plexin-A2 could also influence Purkinje cell and SAC self-avoidance, respectively, through filopodia dynamics (Sun et al. 2013; Matsuoka et al. 2012; Gibson et al. 2014).

Interstitial filopodia also have implications for self-avoidance of 3-dimensional arborizations, which has not yet been definitively demonstrated in mammals. Filopodia bridges that localize γ-Pcdh clusters are observed in hippocampal neural cultures, indicating their prevalence in mammalian dendritogenesis (Fernández-Monreal et al. 2009). Axonal arbor clumping appears in other neural cell types in cPcdh mutants but the underlying repulsive defects are not clear (Mountoufaris et al. 2017; Katori et al. 2017; Prasad and Weiner 2011). An attractive possibility is that the cPcdhs play a universal self-avoidance role through interstitial processes and contact-based interactions that influence local branching decisions.

### Self-contacting filopodia have slower dynamics and require the γ-Pcdhs for contact-induced retraction

A second prominent feature of SAC dendrite self-avoidance is that the self-contacting filopodia population has distinct dynamics. Moreover, these filopodia events are selectively affected by retraction deficits in the absence of the *Pcdhgs*. Self-contacting lifetimes are slower and ∼20-25% of self-contacts are maintained beyond the imaging period. These loops could indicate local decisions for self-contacts to persist and potentially contribute to morphogenetic processes or branch stabilization. By contrast, most non-self-contacting filopodia retract (∼93% across P2-P8), and their motility are unaffected by *Pcdhg* loss. We propose that the self-contacting dynamics reflect SAC dendritic self-recognition events.

The accumulation of self-contacts in *Pcdhg* mutants indicate that the γ-Pcdhs are not required for self-contact formation *per se* but rather regulate a decision to retract. The remaining α-Pcdh and β-Pcdh isoforms might compensate for self-recognition and self-contact in *Pcdhg* SACs, but lack the γ-Pcdhs to induce retraction (**Figure 7**). We have shown that the *Pcdhgs* are essential for SAC self-avoidance, and a single *Pcdhg* isoform rescues self-avoidance (Lefebvre et al. 2012). By contrast, *Pcdha* and *Pcdhb* are dispensable for SAC self-avoidance (Ing-Esteves et al. 2018). However, the more severe self-avoidance defects in double *Pcdha; Pcdhg* mutant SACs suggest more pronounced retraction deficits (Ing-Esteves et al. 2018). Further genetic analyses and live imaging are required to determine how cPcdh diversity or a single γ-Pcdh isoform regulates self-avoidance dynamics.

The excess adhesion but eventual elimination of filopodia loops in *Pcdhg* SACs (compare P8 to P12 in Figure 6B, C) suggest a developmental phase of increased adhesion which might be required for filopodia self-sampling. The adhesive phenotype resembles the excess homotypic fasciculation in other retinal cell types in mouse *Dscam* mutants (Fuerst et al. 2008; Fuerst et al. 2009). In mice, Dscam masks the adhesive properties of other molecules, such as Cadherin members, to limit homophilic fasciculation of neurites belonging to the same cell type (Garrett et al. 2018). Developing SACs co-fasciculate the dendritic processes to form the SAC plexus though the underlying adhesive cues are unknown (Ray et al. 2018; Duan et al. 2018). One possibility is that the γ-Pcdhs selectively mask homotypic adhesion between isoneuronal processes but not between heteroneuronal dendrites. A second possibility is that the heteroneuronal adhesion and the isoneuronal retraction mechanisms work in parallel. In this scenario, self-recognition mediated by homophilic cPcdh isoforms elicits a retraction response.

The γ-Pcdhs likely signal retraction through actin cytoskeletal remodeling but downstream pathways remain elusive. In other contexts, the γ-Pcdhs limit intracellular kinase activities, such as Pyk2/FAK and PKC, to regulate cortical dendrite branching (Suo et al. 2012; Garrett et al. 2012). The α-Pcdhs regulate cytoskeletal dynamics via the WAVE complex to promote cortical neuron migration (Fan et al. 2018). Our approaches to manipulate and image SACs will enable interrogations on self-avoidance signaling. To date, studies linking recognition molecules to filopodia dynamics have advanced mechanistic insight into synaptogenesis and pattern formation but have been limited to a handful of vertebrate cell types (Kayser et al. 2008; Mao et al. 2018; Chen et al. 2010).

How do the γ-Pcdhs selectively respond to self-contacts? The cPcdh self-avoidance model posits that SACs discriminate self from non-self through homophilic interactions between self dendrites expressing the same cPcdh isoforms (Lefebvre et al. 2012). This model is strengthened by recent molecular studies demonstrating the exquisite specificity of cPcdh binding to matching isoforms *in trans* via zipper-like chains (Nicoludis et al. 2015; Rubinstein et al. 2015; Brasch et al. 2019; Goodman et al. 2022; Goodman et al. 2016; Thu et al. 2014). We propose that the longer self-contacting lifetimes reflect such homophilic interactions. The SAC population broadly expresses the *Pcdhas* and *Pcdhgs* (Lefebvre et al. 2012; Ing-Esteves et al. 2018) but isoform expression in individual SACs is unknown. Further investigations of self/non-self discrimination dynamics will require multi-channel imaging to visualize non-self interactions, and analysis of SACs that express the identical or reduced γ-Pcdh isoform diversity (Lefebvre et al. 2012; Garrett et al. 2019).

### γ-Pcdh-dependent self-avoidance is a self-organizing mechanism to produce robust, type-specific morphology

SACs develop a radial dendritic architecture that is conserved across mammalian species and bears little cell-cell variation (Famiglietti 1983; Rodieck 1989; Zhang et al. 2019; Poleg-Polsky et al. 2018). Despite the invariant pattern, SAC dendrites do not grow in a deterministic fashion. By analyzing dendritic growth patterns on the scale of minutes and days, our work reveals the stochastic and self-organizing features of SAC morphogenesis. Recent quantitative studies of *Drosophila da* dendrite morphogenesis showing the important interplay of noisy neurite growth with local repulsive and stabilizing interactions in pattern formation (Ferreira Castro et al. 2020; Palavalli et al. 2021; Shree et al. 2022). Computational modeling demonstrates that stochastic dendrite growth along with negative feedback, such as self-avoidance, recapitulate *in vivo* arborization patterns (Ferreira Castro et al. 2020; Palavalli et al. 2021; Shree et al. 2022). Our quantitative study of SAC dendrite patterning provides a mammalian example of stochastic dendritic exploration coupled with iterative self-avoidance that generate a stereotyped neuronal morphology. Non-deterministic growth guided by local decisions, such as limiting branching or synaptic partnerships, is proposed to be a driver of circuit patterning and wiring specificity (Bassan and Hiesinger 2015; Wit and Hiesinger 2022). The expansion of methods to quantify branching morphogenesis and dynamics (Hogg et al. 2021; Wang and Lefebvre 2022) will undoubtedly present new avenues to study self-organizing features and other patterning mechanisms that specify diverse neuronal morphologies.

## METHODS

### Mouse strains

Mouse lines used in this study have been previously described. The conditional *Pcdhg*^*fcon3*^ allele contains LoxP sequences that flank the third constant exon (Lefebvre et al. 2008; Prasad et al. 2008). Mice with SAC-specific *Pcdhg* deletions (denoted by *Pcdh SACko*) were obtained by crossing the *Pcdhg*^*fcon3*^ line to *ChAT-IRES-Cre* mice (B6;129S6-*Chat*^*tm2(cre)Lowl*^*/*J; RRID:IMSR_JAX:006410; Rossi et al. 2011). The Cre reporter *ROSA*^*mT/mG*^ line was obtained from Jackson Laboratories (Gt*(ROSA)26Sor*^*tm4(ACTB-tdTomato,-EGFP)Luo*^*/*J; RRIP:IMSR_JAX:007576; Muzumdar et al. 2007). Mice were maintained on a C57/B6J background. All experiments were carried out in accordance with the Canadian Council on Animal Care guidelines for use of animal in research and laboratory animal care under protocols approved by the Centre for Phenogenomics Animal Care Committee (Toronto, Canada) and the Laboratory Animal Services Animal Care Committee at the Hospital for Sick Children (Toronto, Canada).

### Transgenic and AAV Labeling of Neurons

Sparse labeling of SACs at P0-P4 was obtained by membrane-localized eGFP expression from *ChAT-IRES-Cre; ROSA*^*mT/mG*^ mice. Sparse labeling of SACs for P6 and beyond was obtained by injection of Brainbow AAVs encoding farnesylated fluorescent proteins in P0 retina, as previously described (Ing-Esteves and Lefebvre 2021). In all experiments, displaced SACs residing in the ganglion cell layer were selected for imaging. Recombinant AAV9-hEF1a-LoxP-TagBFP-LoxP-eYFP-LoxP-WPRE-hGH-InvBYF and AAV9-hEF1a-LoxP-mCherry-LoxP-mTFP-LoxP-WPRE-hGH-InvCheTF viruses were a gift from Dawen Cai & Joshua Sanes (Cai et al. 2013) and obtained from the University of Pennsylvania Vector Core (Addgene viral preps # 45185-AAV9; # 45186-AAV9). A total of ∼9 × 10^8^ to 2 × 10^9^ viral genome particles of each vector (∼0.25 µL of 3-4 × 10^12^ GC per mL dilution in sterile PBS with 0.02% Fast Green FCF dye, pH = 7.4) were injected intravitreally into retinas of *Pcdhg*^*fcon3*^; *ChAT-IRES-Cre* P0 animals using a Pneumatic PicoPump (World Precision Instruments).

### Imaging of retinal explants

Detailed protocols for live-imaging of *ex vivo* retinal explants by scanning confocal microscopy and analyses has been previously described (Ing-Esteves and Lefebvre 2021). Briefly, retinas were dissected in oxygenated artificial cerebrospinal fluid (aCSF, Williams et al. 2013) and mounted flat onto mixed cellulose ester (MCE) membrane filter discs (Millipore HABG01 300). Live retinal flat mount preparations were perfused with heated and oxygenated aCSF (temperature 32-34 °C, flow rate 1 mL/min). Samples were imaged with the following parameters: an upright Leica SP8 with Super z-Galvo stage, 488-nm or 552-nm illumination at 1% to 4% laser power, a 25x water immersion objective with an 0.95 numerical aperture (Leica HC FLUOTAR L 25x/0.95 W VISIR lens; XY resolution of 256.84nm), and LASX software. Full-frame movies (1024 by 1024 pixels) containing 10-15μm volumes with step size 0.4-1μm were collected every 2-2.25 minutes. Imaging durations were limited to 30-60min to prevent excess heating or phototoxicity. Static images were either obtained with the 25x or 63x glycerol immersion lens with 1.3 NA (Leica HC PL APO 63X/1.3 GLYC CORR CS2; XY resolution of 187.69nm).

### Image Processing and Analysis

Movies were curated, and image pre-processing and analysis was performed using NIH ImageJ/Fiji software (Schindelin et al. 2012) as described previously (Ing-Esteves and Lefebvre 2021). Resolution enhancement of 3D volumes by deconvolution was performed by generating a theoretical point spread function (PSF) using the ImageJ plugin ‘Diffraction PSF 3D’ (https://imagej.net/plugins/diffraction-psf-3d), and applying it to deconvolve each time point of the image series. For morphometric quantifications of static images, deconvolved 3D image volumes at single time points were selected, and the region of interest (ROI) encompassed the entire arbor. For dynamic filopodia analyses, Z-dimension maximum intensity projections of imaging volumes were created for each deconvolved time point to create a 2D time series. Registration of 2D time points was performed with the ImageJ plugin StackReg (ImageJ: https://imagej.net/plugins/stackreg). ROIs of 15 × 15μm or 20 × 20μm were selected in each quadrant of the arbor. For the SAC arbors at P2-4, ROIs were adjacent to the soma and encompassed both proximal and distal arbor regions. For P6-8 arbors, ROIs were selected at both proximal and distal regions of the arbor. On average, 1-3 cells were imaged per animal, and 2-4 ROIs were analyzed per cell. Video durations ranged from 30-65 minutes and quantifications were normalized to a 20μm x 20μm region observed for 1 hour (*i*.*e*. count/400μm^2^•hr).

Filopodia self-contacts and non-contacting events were manually annotated and quantified. Filopodia self-contacts were called by a continuous fluorescent signal forming a loop, and were manually verified against the corresponding 3D image series. Filopodia lifetime is the interval between filopodia emergence and retraction, and calculated as the number of frames during which the filopodia are present, minus 1, then multiplied by the time-step for the movie.

For terminal branch point quantifications, endpoint processes at distal-most locations of the arbor were selected, and included branch points and filopodia not connected to any other point on the dendrite tree. To determine the asymmetry index, we calculated the radial variance of pixel content of Z-projections of SAC images that were converted to a binary threshold and analyzed using the ImageJ plugin “Azimuthal Average” with a bin size of 8 (Ing-Esteves et al. 2018). The standard deviation between bins was calculated and then normalized against average pixel content value of the entire SAC arbor to produce a unitless radial asymmetry index. Batch calculations for 5-6 neurons for each age and genotype were performed using a Python3 script. Circularity and convex hull were measured using ImageJ.

### Experimental design and statistical analysis

Morphometric and dynamic filopodia quantifications were performed in at least three animals for each genotype and from multiple litters. Cre-positive WT or heterozygous *Pcdhg*^*fcon3*^ littermates served as controls. Both male and female animals were analyzed. The analyst was blinded to the genotypes for all analyses. Statistical analyses were performed in Python3 using either the Pingouin 0.3.12 (https://pingouin-stats.org/) or SciPy 1.5.4 (https://scipy.org/) statistical packages. Data are presented as means with standard deviation unless otherwise stated.

Means of two groups, including those of control to *Pcdh SACko* cells at specific ages, were compared using two-sample, two-tailed Student’s t-test for normally distributed samples, or two-tailed Mann Whitney non-parametric test. One-way or two-factor ANOVA followed by post-hoc pairwise analyses were used for comparisons of means across ages and/or genotype. 1-sample Student’s t-test was used to determine if a single population mean was significantly different from the null hypothesis of 50%. Exact p-values are reported unless the values are <0.0001.

### Histology

Anesthetized mice were sacrificed by decapitation and enucleated immediately in PBS. Eye cups were dissected on ice in PBS and retinas fixed in 4% PFA at 4°C for 6–12 h. Whole-mount preparations and cryosections of retinas were prepared as described previously (Lefebvre et al. 2012; Ing-Esteves et al. 2018). Whole retinas were incubated successively with blocking solution (0.3-0.5% Triton-X, 8% normal goat serum in PBS) and then with primary antibodies for 2 days at 4°C. After washing, retinas were incubated for 3 hours at room temperature with Alexa Fluor-conjugated secondary antibodies (1:1,000; Thermo Fisher Scientific or Jackson ImmunoResearch). Whole retinas were flattened with photoreceptor side down onto nitrocellulose filters. Isolated SAC image was obtained from SAC cultures described in (Lefebvre et al. 2012). Primary antibodies used for this study were as follows: chick anti-GFP (1:2000, Aves Laboratories, GFP-1010, RRID:AB_2307313); goat anti-choline acetyltransferase(1:250, Millipore, AB144P, RRID:AB_2079751); goat anti-vesicular acetylcholine transporter (1:1000, Millipore, ABN100, RRID:AB_2630394); mouse anti-syntaxin HPC1 clone (1:1000; Sigma SAB4200841).

## ACKNOWLEDGEMENTS

This work was supported by an NSERC Discovery grant (RGPIN-2016-06128); a Canada Research Chair (Tier 2), and a Sloan Fellowship in Neuroscience (FG-2015-65234) to J.L.L. S. Ing-Esteves was supported by the Vision Science Research Program and an NSERC Postgraduate Scholarships-Doctoral Program.

## AUTHOR CONTRIBUTIONS

S. I-E. and J.L.L. conceptualized the project and designed experiments. S. I-E. performed experiments and computational analyses. S. I-E. and J.L.L. wrote the manuscript.

